# Distinguishing word identity and sequence context in DNA language models

**DOI:** 10.1101/2023.07.11.548593

**Authors:** Melissa Sanabria, Jonas Hirsch, Anna R. Poetsch

## Abstract

Transformer-based large language models (LLMs) are very suited for biological sequence data, because the structure of protein and nucleic acid sequences show many analogies to natural language. Complex relationships in biological sequence can be learned, although there may not be a clear concept of words, because they can be generated through tokenization. Training is subsequently performed for masked token prediction. With this strategy, the models learn both the token sequence identity and a larger sequence context. We developed a framework to interrogate what a model learns, which is both relevant for the interpretability of the model and to evaluate its potential for specific tasks.

We used a DNA language model, which is trained on the human reference genome with a Bidirectional Encoder Representations from Transformers (BERT) model. In this model, tokens are defined with overlapping k-mers. To gain insight into the model’s learning, we interrogated how the model performs predictions, extracted token embeddings, and defined a fine-tuning benchmarking task to predict the next tokens of different sizes without overlaps. This task is very suited to evaluate different pretrained DNA language models, also called foundation models, since it does not interrogate specific genome biology, does not depend on the tokenization strategy, the size of the vocabulary, the dictionary, or the number of parameters used to train the model. Lastly, the task performs without leakage of information from token identity into the prediction task, which makes it particularly useful to evaluate the learning of sequence context.

Through this assessment we discovered that the model with overlapping k-mers struggles to learn larger sequence context. Instead, the learned embeddings largely represent token sequence. Still, good performance is achieved for genome biology inspired fine-tuning tasks. Models with overlapping tokens may be used for tasks where a larger sequence context is of less relevance, but the token sequence directly represents the desired learning feature. This emphasizes the need to interrogate knowledge representation in biological large language models. Transparency is particularly important for biomedical use cases and an understanding of what the models are learning can be used to match the model to the desired task.

## Introduction

With the developments of novel deep learning strategies, there has been rapid progress in how we can understand and generate language. Models for Natural Language Processing (NLP) are frequently based on transformer architectures^1^ and thus do not only show unprecedented performance, but through the transparent nature of these architectures, it can be extracted what and why the models learn. Together this makes them particularly attractive for non-language tasks that resemble language, such as the code-breaking of sequence data in biology, particularly protein sequence and nucleic acids. We know that there is structure in these data, but of much of the sequence we understand very little. In DNA we have a code for translation into proteins, sequence-dependent gene regulation and regulation of retrotransposons and repetitive sequence. Also, chromatin architecture, DNA replication, and genome stability itself, all depend partially on sequence, but how these codes interact and function in detail has been out of reach until now. To dissect the relationships and rules, machine learning and deep learning have been applied to grasp specific genome features with various architectures from simple neural networks and convolutional neural networks^2^ to combining those with transformers^3^. This model, called Enformer, allows a wide learning range of up to 100 kb length and unprecedented performance of gene expression prediction. These models have however been engineered for specific tasks and therefore focus on one aspect of the genetic code. In NLP, pretrained large language models (LLMs), like GPT-3^4^ and successors have changed how such models see language, and they have been built as foundation models that can be fine-tuned for a variety of downstream tasks.

Recently, LLMs have been adapted to DNA sequence with the aim of building foundation DNA language models that learn information resembling grammar and syntax. Several foundation DNA language models have now been made available, specifically Nucleotide Transformer^5^ and DNABERT^6^. However, DNA sequence differs from natural language, most prominently by “words” not being naturally defined in DNA. Still, they can be artificially generated through tokenization, i.e. by defining groups of nucleotides as separate tokens. Tokens are embedded, which has the consequence that information on the original underlying sequence is initially not retained. For generating the tokens, the models use different strategies. For Nucleotide Transformer, consecutive 6mers are used, complemented by monomers around edges and unknown sequence^5^. DNABERT on the other hand trains on overlapping k-mers^6^. Both models are based on Bidirectional Encoder Representations from Transformers (BERT)^7^ architectures. With the learning process through the transformer architecture, the model itself is training larger sequence context by updating its attention. Also the embedding is updated, which gives the tokens order relative to each other. From this it can be extracted and analyzed how the model sees the tokens.

Foundation models can be finetuned for a variety of tasks, including classification, regression, and generative problems. This way, functional elements like promoters can be detected, as well as splice sites, transcription factor binding sites, enhancer function and/or gene expression. Different base models may have different suitability for diverse tasks, because for some tasks the identity of individual tokens is of high importance for the training, whereas other tasks rather require learning of larger sequence contexts.

On the example of DNABERT, we have built a framework to investigate the learning of foundation DNA language models using specialized fine-tuning tasks, extraction of predictions, and embeddings. We thus could show that a large language model that is trained with overlapping tokens predominantly learns token identity. While larger context learning is limited, the model is particularly suited for fine-tuning tasks that require sequence knowledge.

## Results

To build a framework to extract information content from foundation DNA language models, we used DNABERT^6^, a transformer model^1^ with a Bidirectional Encoder Representations from Transformers (BERT)^7^ architecture (Fig.1). The model uses the hg19 human reference genome tokenized to 4mers, 5mers, and 6mers. It is trained for cross-entropy loss by spligng the genome into training, validation and test windows to predict masked tokens of the same number as token length. Through the design of using tokens from overlapping k-mers, unmasked sequence partially shares sequence with the masked tokens. The central nucleotide of the combined masked tokens is the only nucleotide that is completely masked.

**Figure 1.**
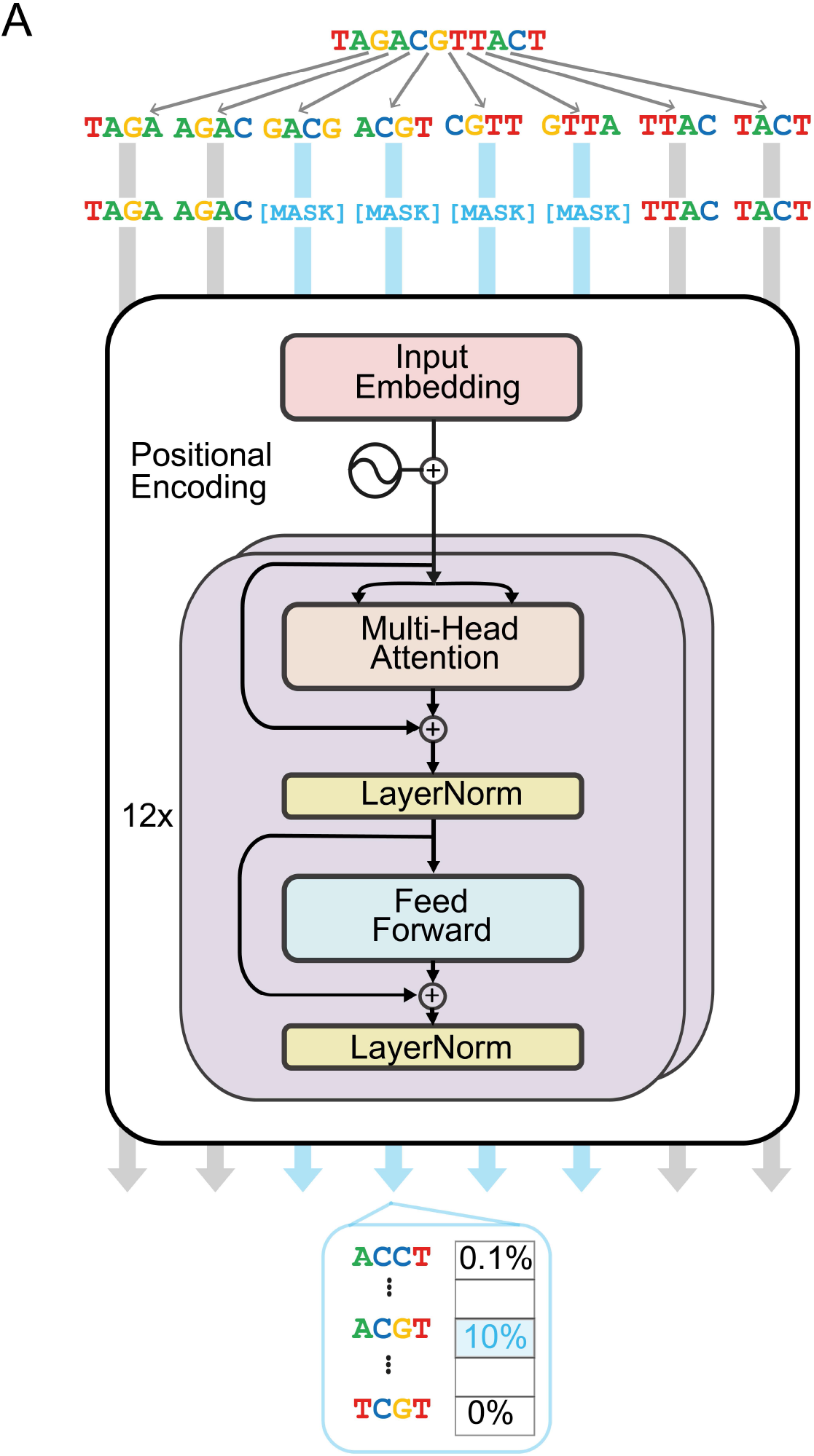
The model architecture of DNABERT. The model is a BERT architecture with 12 transformer blocks. It splits a given sequence into tokens, which are overlapping k-mers, 4, 5, or 6 nucleotides in length. The model is embedding the tokens and is trained with cross-entropy loss to predict the masked tokens and updates the embedding while training. The number of masked tokens is equivalent to token size, which leaves the central nucleotide free from overlap. The model outputs probabilities of token identities of the masked tokens.

### Prediction performance for masked k-mers with overlaps

We investigated on some individual random representative examples how the training strategy leads to prediction of probabilities of tokens (Fig.2A). Although the model initially does not include knowledge of token identity, it becomes clear that the training leads to the model restricting its choice to four k-mers, where the central nucleotide of the masked tokens is predicted with variable probabilities. The most, second-most, third-most, and fourth-most likely tokens are each respectively assigned similar probabilities throughout the masks, even in samples that show poor performance. All other tokens are predicted with close to zero probabilities.

**Figure 2.**
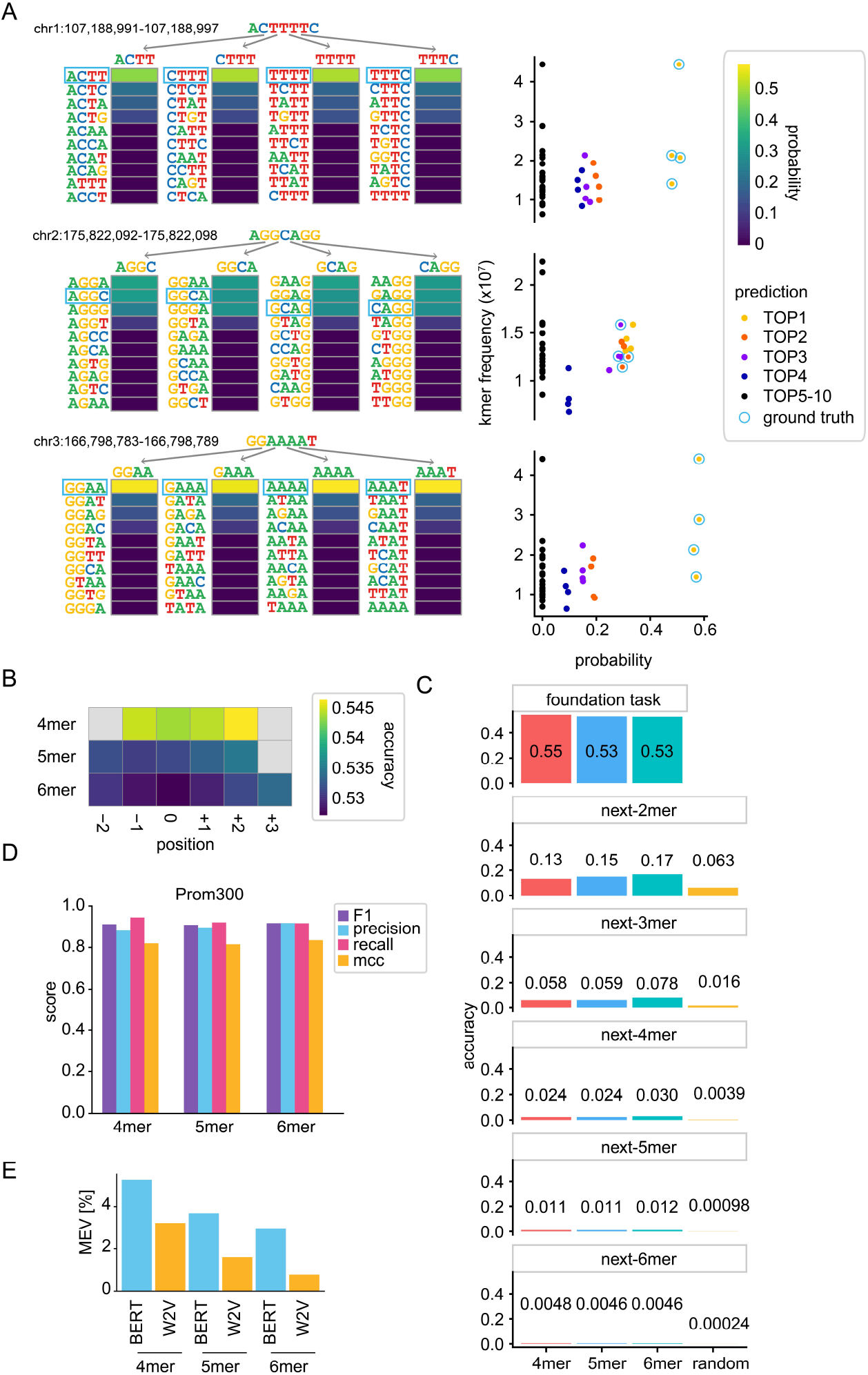
Representation and context learning. **(A)** Token prediction probabilities on 3 random examples of 7 consecutive nucleotides in the genome with the 4mer DNABERT model. Ground truth is framed in blue. On the left, probabilities are given per position, on the right in comparison with k-mer frequency in the genome and color-coded by prediction rank per 4mer. **(B)** Average accuracy of token prediction per k-mer relative to the central token position. **(C)** Accuracy of token prediction as a fine-tuning task for the foundation models using prediction of 2 to 6 nucleotide long next-k-mers as readout. Comparison to random k-mer accuracy and the accuracy of the foundation models, i.e. average token prediction of the masked tokens of the respective k-mer model **(D)** Performance metrics for the Prom300 finetuning task to identify promoter structures by classifying real promoters versus promoters with shuffled nucleotides (left). Metrics are given as F1 score, precision, recall, and Mathews Correlation Coefficient (MCC). **(E)** Maximum explainable variance through the embedding of the BERT model versus Word2Vec (W2V) static embedding of the k-mers.

On average, the training task is performed with an accuracy of 0.544, 0.532, and 0.527 for the central 4mers, 5mers, and 6mers in the test set (Fig2B). For all k-mer models there is slightly improved performance for the outer masked tokens, so that the average performance is 0.545, 0.533, and 0.530 for 4mers, 5mers, and 6mers respectively (Fig2C). These results indicate that the model learns sequence identity of tokens, which raises the question of how much sequence context it learns in addition and how much this is required for biological fine-tuning tasks. For many biological questions, a larger sequence context is of critical importance and this is a main reason why large language models can be so powerful on biological data. However, from a biological task it is difficult to differentiate whether the token sequence itself or larger sequence content is the relevant learning feature, because many tasks are dependent on both. For example, transcription factor binding frequently requires a motif, but also other transcription factor binding sites nearby and/or specific physicochemical properties of the surrounding DNA^8^. Thus, evaluating foundation models only on biological fine-tuning tasks cannot sufficiently differentiate between these different learning features.

### A universal fine-tuning task for foundation DNA language models

We aimed for a fine-tuning task to use as a general benchmark for learning of sequence context in DNA language models. Such a task needs to fulfil three criteria. First, it needs to be independent from the biology we are interested in, second it needs to be a task that can be used to compare different models and thus does not depend on tokenization strategy, the size of the vocabulary or the dictionary, and the number of parameters the model was trained on. Third, the task needs to require sequence context for performance without leakage of information from token identity into the prediction task. These criteria are all fulfilled with a fine-tuning task for next-token prediction. We set up 5 models that are fine-tuned to predict next-2mers to next-6mers, irrespective of whether the respective next-k-mer size was part of the tokenization strategy in the foundation model. When applying this task to DNABERT (Fig 2C), performance drops to 0.024, 0.024, and 0.030 for predicting next-4mers with the models trained for overlapping 4mers, 5mers, and 6mers respectively, which compares to an accuracy of 0.004 for a random next-4mer. Interestingly, if the token size of the foundation model matches the token size of the fine-tuning task, performance does not seem to improve over different token sizes. It can thus be concluded that the model learns very little context that is of help to predict sequence which is not overlapping with the training task.

### Promoter identity training learns sufficiently from sequence identity

Since DNABERT still shows good performance with biological training tasks^6^, we interrogated further the Prom300 fine-tuning task, which is a classification task for promoter identity with promoters of 300 base pairs length. The promoter sequence is mutated and DNABERT is trained to distinguish real promoters from mutated promoters. It does this with an F1 score of 0.965^6^. Since mutation may change the sequence content of the promoters, we adjusted the task to rather shuffle the promoter sequence on the nucleotide level (Fig 2D). Also with this adjusted task, DNABERT discriminates promoters from shuffled promoters with an f1 score of 0.91, 0.91, and 0.92 for 4mers, 5mers, and 6mers respectively. Thus, this fine-tuning task does indeed benefit from the sequence learning of the tokens. It can be expected that also other biological fine-tuning tasks may similarly benefit from this learning feature of the foundation model.

### Sequence context gets embedded in the tokens

Another measure for learning of grammatical context in a large language model is to investigate how contextual the contextualised k-mer representations in the embedding for the model really are^9^. This can be quantified through the maximum explainable variance (MEV), the variance explained on the first principal component when applying a principal component analysis (PCA) on average trained embedding of the tokens. A small variance explained indicates learning of context rather than word representation. For comparison to the BERT model, we used an algorithm that estimates word associations in vector space through static embeddings of context, Word2Vec (W2V)^10^. Since W2V is an algorithm that through the static embeddings cannot represent lexical ambiguity, it would be expected to learn little individual context and thus learn a larger proportion of word identity leading to a higher variance explained. Surprisingly, however, the BERT models all lead to a MEV that is higher than the equivalent MEV for W2V (Fig 2E). We thus applied a non-linear dimensionality reduction algorithm Uniform Manifold Approximation and Projection (UMAP)^11^ to investigate the learned embedding for non-linear relationships (Fig3). Analyzing the clustering of the embedding of the 4mer model (Fig3A), it becomes clear that BERT groups the 256 tokens into 16 clusters, four groups of four. The subclusters have the central dinucleotide in common, whereas clusters of similar pyrimidine-to-purine patterns cluster more closely. Investigating the W2V embedding, the clustering is visible, but not as apparent as with the BERT embedding. For the 5mer model, the difference of the clustering between BERT embeddings and W2V embeddings becomes more apparent (Fig.3B). The BERT embedding groups the tokens roughly into 16 groups of 4, dependent on the pyrimidine-purine balance of the central trinucleotide. W2V forms some clusters, but does not seem to be able to distinguish all token groups. For the 4,096 6mers (Fig 3C), the BERT embedding also generates 4 groups of 4, whereas W2V forms some small dense clusters, but does not differentiate between all token groups. It can thus be concluded that the embedding largely represents the sequence identity of the learned tokens. Given that this clustering becomes only clearly apparent with a non-linear method, rather than PCA (data not shown), the maximum explainable variance is probably still an underestimate, since it does not account for variance to be explainable in a non-linear space.

**Figure 3.**
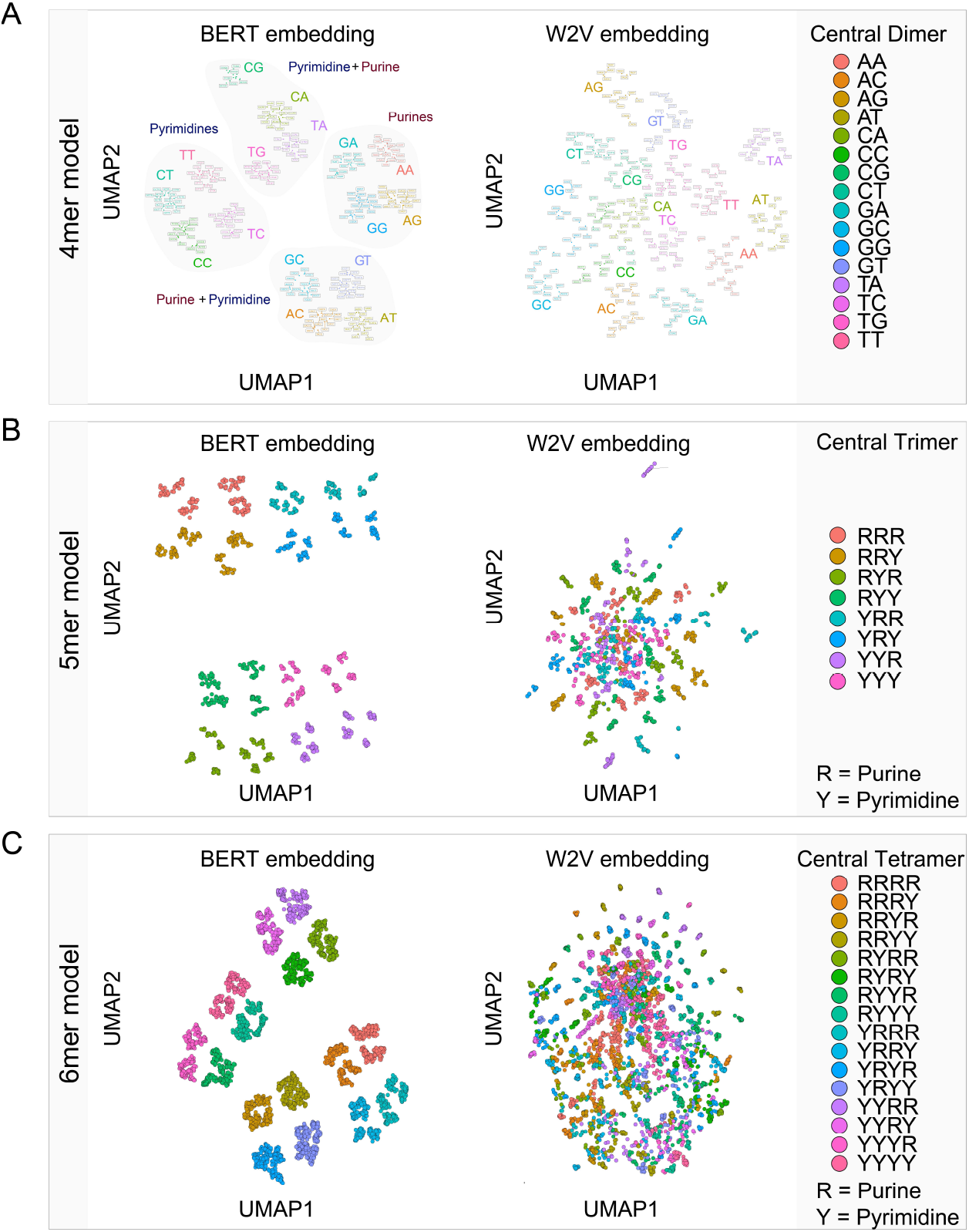
Extracting representations from contextualized embeddings. Uniform Manifold Approximation and Projections (UMAP) of k-mer embeddings comparing BERT embeddings versus Word2Vec (W2V) embedding applied to **(A)** 4mers with highlighting the central dimer and its pyrimidine-purine balance, **(B)** 5mers with the pyrimidine-purine balance of the central trimer, and **(C)** 6mers with the pyrimidine-purine balance of the central tetramer.

## Discussion

What DNA language models learn is of crucial importance to allow interpretation of the tasks they perform. We have built a framework to differentiate context learning from token representation through its sequence to investigate the learning of DNABERT. Using extraction of predicted probabilities, we discovered that the model largely decides between four k-mers, only different by the nucleotide in the center of the masked tokens. Overall average accuracy of token prediction is above 50%. To compare the models on a unifying task, but not biased by a specific biological question and with a focus on context rather than sequence identity, we developed a task of next-token prediction that can be used to generally compare foundation DNA language models. In this task, DNABERT struggled to predict next-k-mers of the same size that it managed to predict when masked. While performance works well for promoter identification, evaluation for contextualized learning with maximum explainable variance also showed that average embedding of the tokens explains more maximum variance than the static W2V embedding. UMAP dimensionality reduction of the embedding revealed that indeed in the embedding the tokens are represented largely as the spelled-out sequence, rather than in context.

Models that use overlapping tokens are thus probably not suitable for tasks that require a larger sequence context, such as protein-DNA binding, which is mediated beyond the DNA binding motif, or tasks for combinatorial genotype-to-phenotype predictions. However, learning of sequence is also of importance for many biological tasks, which explains the good performance of the model in several biological finetuning tasks from the original study^6^, especially when the identity or the specific positions of the tokens matter (e.g. in a promoter). However, what this study exemplifies is that for using such models as widespread in genetics as can currently be anticipated, it is very important that we understand what we are training for and what we are learning. Similar to NLP we need to build tools and frameworks to test, whether the models are really learning what we designed them for. In genomics this also becomes increasingly important, since there are also risks of increasing and perpetuating biases, such as biases from the use of reference genomes, incomplete or false genomics data. Unlike a natural language generative AI, where racist and sexist training biases become visible in the output, we are aiming for unknown discoveries in genomics. Therefore, issues in the training process do not directly become apparent but may have substantial consequences for downstream performance, accuracy of research data, and may especially harbor risks for patients, when used for biomedical research.

## Methods

The final source code and fine-tuned models will be made available at https://zenodo.org

### DNA Language Model Architecture

We train with the *Homo sapiens* (human) genome assembly GRCh37 (hg19), only taking into account the sequences that contain A,C,G and T. We use the pre-trained models and code of the DNA language model DNABERT^6^, provided by the authors (https://github.com/jerryji1993/DNABERT; June 2023). DNABERT is based on a Bidirectional Encoder Representations from Transformers (BERT)^7^ model that takes as input tokenized sequences of up to 510 tokens long. Tokenization is performed with overlapping k-mers, 4, 5 and 6 nucleotides in size, selected based on the performance metrics in the original study. The vocabularies consist of all the permutations of k consecutive nucleotides (i.e. 256, 1024 and 4096 respectively) as well as five special tokens: CLS, PAD, UNK, SEP and MASK. CLS represents the classification token, PAD is used for padding the right side of the sequence in case it is shorter than the maximum input length of the model, UNK is for sequences of nucleotides that do not belong to the vocabulary, SEP is used to indicate the end of a sequence and MASK represents the masked tokens.

### Masked token prediction

To extract what the model has learned, we extract what the model predicts over the mask. Each chromosome is split into sub-sequences. The length of each sub-sequence varies between 20 and 510 tokens. Specifically, with a 50% probability, the length of a sub-sequence is 510. With another 50% probability, its length is a random integer between 20 and 510. Then 20% of the sub-sequences are taken as the dataset for this task, which is around one million samples.

For each of the samples we randomly choose a token and mask tokens of equivalent numbers to the size of the k-mer using the following pattern; 4mer: −1,0,1,2; 5mer: −2, −1,0, 1, 2; 6mer: −2, −1, 0,1, 2, 3. Position 0 represents the chosen token, “-” represents the tokens previous to the central token and “+” the following tokens. This way the central nucleotide of the mask does not overlap with any token outside the mask

### Next k-mer prediction

To build a fine-tuning task that allows to compare different foundation models that relies on context learning, and is not dependent on the biological question, we established next-k-mer prediction. We take the pre-trained language models (4mer, 5mer and 6mer) and fine tune every model to predict the next k-mer, where k is 2, 3, 4, 5 and 6.

To create the data for this task, chromosome 21 is split into sequences of 510 nucleotides. We keep the first 56 nucleotides of each sequence. These sequences are randomly shuffled. Finally, the dataset is composed of 500,000 sequences, where 80% of them are for training and 20% for testing.

The samples are defined as the first 50 nucleotides of each sequence. For the labels, we take the k (2, 3, 4, 5 and 6) nucleotides that follow the 50 nucleotides. The next-kmer model will have 4^k different classes, i.e., 16, 64, 256, 1024 and 4096, respectively, which are all the permutations of k nucleotides.

The models are trained with cross entropy loss on the prediction of the next-k-mer using Adam optimizer with a learning rate of 10^-6, epsilon of 10^-8, and beta of 0,99. The model accepts a maximum input length of 50 tokens. The dropout probability of the classification layer is 0.5. We use batch size of 64, and train for 150 iterations.

### Promoter Identification

The Prom300 task was adapted with some minor modifications from Ji et al.^6^. The modification was made in regards to the disruption of sequence. In short, we use the human data (hg19) from the Eukaryotic Promoter Database (https://epd.ep?.ch/human/human_database.php?db=human) to obtain annotation of 30,000 intact promoter sequences, which we define as 300bp long ranges from −249 to +50 bp around the Transcriptional Start Site (TSS). For the definition of non-promoter samples, we apply a shuffling strategy of nucleotides rather than mutation to prevent changes in the nucleotide composition of the sequence. We divide the sequence into 20 parts of equal size and then shuffle 15. The sequences are tokenized according to each model, i.e. divided in overlapping kmers. For the prediction, we add a classification layer with one neuron. The model is trained with cross entropy loss, using an Adam optimizer with a learning rate of 10^- 6, an epsilon of 10^-8, and beta of 0,99. The model accepts a maximum input length of 50. We use batch size of 64, and train for 10 epochs.

### Word2Vec

For comparison of token embeddings, we use Word2Vec, a static word embedding tool^10^ that maps each word to a single vector. In general, this mapping function does not account for lexical ambiguity, which means that identical letter sequences can have multiple interpretations or different grammatical roles. We implemented Word2vec with a continuous bag-of-words (CBOW) approach for learning representations of words, which uses the surrounding words in the sentence to predict the middle word. The context includes a window of 5 words with the current word in the center. This architecture is referred to as a bag-of-words model because it does not consider the order of words in the context.

To generate the Word2Vec (W2V) embeddings, first each chromosome is split into sub-sequences. The length of each sub-sequence varies between 20 and 510 tokens. Specifically, with a 50% probability, the length of a sub-sequence is 510. With another 50% probability, its length is a random integer between 20 and 510. Then 300,000 of the sub-sequences are randomly chosen as the dataset for this task. We tokenize each sequence with overlapping tokens, and create three datasets, one for each kmer (4mer, 5mer and 6mer). We use the Word2Vec module of Gensim (https://radimrehurek.com/gensim/models/word2vec.html), with the following parameters: min_count = 1, vector_size = 768, window = 5.

### Model Embedding

Unlike static word embeddings, dynamic word embeddings aim at capturing word semantics in different contexts to address issues like the context-dependent nature of words. We obtain a summarized version of the contextualized word representations that is the token embedding of the BERT model. To obtain the token embeddings of the model, we extract from DNABERT^6^ the weights of the layer word_embeddings for each k-mer model.

### Dimensionality reduction and maximum explainable variance

Both W2V and DNA Language Model embeddings are represented as vectors of size 768. Average distances between tokens are thus interrogated through the dimensionality reduction algorithms Principal Component Analysis (PCA) and Uniform Manifold Approximation and Projections (UMAP) in R with the packages ‘stats’ (4.2.1) and ‘UMAP’ (0.2.10.0), respectively. As a measure of context learning, Maximum Explainable Variance (MEV)^9^ was extracted as the variance explained by the first principal component.

## Author contributions

ARP has conceptualized the study, MS and JH applied the models and implemented the fine-tuning tasks. All authors designed the fine-tuning tasks and analyzed the data. MS and ARP wrote the manuscript with input from JH.

## Acknowledgement

This work was supported by the Center for Scalable data analytics and artificial intelligence (Scads.AI) Dresden-Leipzig. **ARP** was supported by the Mildred Scheel Early Career Center Dresden P2, funded by the German Cancer Aid. **MS** was supported by a TU Dresden Junior Fellowship and a Maria Reiche Postdoctoral fellowship of TU Dresden. **JH** was supported by the TU Dresden program ‘FOSTER – Funds for Student Research’.

